# Oxygen radicals and cytoplasm zoning in growing lily pollen tubes

**DOI:** 10.1101/2020.05.01.069807

**Authors:** Alexandra Podolyan, Oksana Luneva, Maria Breygina

## Abstract

Recently redox-regulation of tip growth has been extensively studied, but differential sensitivity of growing cells to particular ROS and their subcellular localization is still unclear. Here we used specific dyes to provide mapping of H_2_O_2_ and O^•^_2_^−^ in short and long pollen tubes. We found apical accumulation of H_2_O_2_ and H_2_O_2_–producing organelles in the shank that were not colocalized with O^•^_2_^−^-producing mitochondria. Differential modulation of ROS content of the germination medium affected both growth speed and pollen tube morphology. Oxygen radicals affected ionic zoning: membrane potential and pH gradients. OH• caused depolarization all along the tube while O^•^_2_^−^ provoked hyperpolarization and cytoplasm alkalinization. O^•^_2_^−^ accelerated growth and reduced tube diameter, indicating that this ROS can be considered as pollen tube growth stimulator along with H_2_O_2_. Serious structural disturbances were observed upon exposure to OH• and H_2_O_2_ and O^•^_2_^−^ quencher MnTMPP: pollen tube growth slowed down and ballooned tips formed in both cases, but in the presence of OH• membrane transport and organelle distribution was affected as well. OH•, thus, can be considered as a negative influence on pollen tubes which, presumably, have mechanisms for leveling it. The assumption was confirmed by EPR spectroscopy: pollen tubes actively reduce OH• content in the incubation medium.

## Introduction

Pollen tube growth is designed to deliver sperms to the ovule, which ensures successful transmission of hereditary information and the alteration of generations. The growing tube is characterized by important structural features typical for cells with polar growth, such as cytoplasm zoning. The key compartments: apical, subapical and distal – differ in cytological parameters: composition of organelles, trajectory of cytoplasm movement and cytoskeleton structure (Cheung and Wu 2008). The distribution of inorganic ions is also strictly subjected to the principle of zoning, including gradient distribution of ionic currents through the plasma membrane as well as free inorganic ions in the cytoplasm (Michard et al. 2017; Feijó and Wudick 2018; Zheng et al. 2019). The latter is maintained due to the uneven allocation of ion transport systems and their activity – which also results in the gradient distribution of membrane potential values, which has been to date revealed in 3 species, including lily (Breygina et al. 2010; Maksimov et al. 2018; Podolyan et al. 2019).

Recent studies of pollen tube growth have been attributed to internal and external signals affecting cytoplasm structural and functional as well as non-uniformly distributed ion currents. A number of data indicate the great importance of redox-regulation: ROS are produced on stigmas of various plants species during the pollination period (McInnis et al. 2006; Zafra et al. 2010) and also accumulate in other parts of the pistil, including the ovule (Duan et al. 2014). At the same time, in NADPH oxidase mutants pollen tubes exhibit anomalies, including impaired intracellular CA^2+^ dynamics and a decrease of the inward K^+^ current (Lassig et al. 2014), while transient suppression of this enzyme caused general growth inhibition (Potocký et al. 2007; Jimenez-Quesada et al. 2019). Another source for ROS proposed more recently is polyamine oxidase (PAO), which uses polyamines as substrate to produce hydrogen peroxide (Wu et al. 2010); polyamines are believed to be loaded into vesicles and transported to the tube apex, which was demonstrated to be crucial for pollen tube growth (Do et al. 2019). These data suggest that fertilization requires both exogenous ROS produced by female sporophyte tissues and endogenous ROS produced by the male gametophyte.

The spatial distribution of endogenous ROS in pollen tubes has been mostly visualized by non-specific staining; tip-localized ROS were found in *Pyrus, Cupressus* and *Picea* pollen tubes (Liu et al. 2009; Aloisi et al. 2015; Pasqualini et al. 2015). Recently H_2_O_2_ accumulation was reported in short *Arabidopsis* tubes (Do et al. 2019). Here we use a combination of dyes that had been applied previously to study *Picea* pollen tubes (Maksimov et al. 2018), which exhibit multiple differences from angiosperms, including redox-regulation and cytoplasm flow.

Model experiments on protoplasts with exogenous ROS highlighted several aspects of pollen physiology related to redox-signaling, the main of which is ion homeostasis, as ion transport across the membrane is regulated by H_2_O_2_ (Breygina et al. 2016; Maksimov et al. 2016). As H_2_O_2_ is the most stable ROS, all studies *in vitro* were confined to H_2_O_2_, which was shown to have multiple effects in pollen tubes, both angiosperm (Podolyan et al. 2019) and gymnosperm (Maksimov et al. 2018). Studies on somatic plant cells that have been exposed to stress lead to the opinion that H_2_O_2_ doesn’t affect “targets” that had been attributed to this ROS but rather binds to transition metal binding sites converting it to OH^•^ (Demidchik 2015). To distinguish between the effects of two ROS we need to compare them directly. Also the formation of OH• could be detected by EPR spectroscopy. To date, none of these tests still had been carried out. Inhibitory analysis indicated the possible involvement of other ROS, namely, OH• and O^•^_2_^−^, in pollen germination (Smirnova et al. 2013), but the effects of exogenous radicals have not been studied to date, despite their possible involvement in growth regulation *in vivo*. Here we study the regulation of pollen tube growth, cytoplasm zoning, membrane potential gradient and pH gradient by exogenous free oxygen radicals: OH^•^ and O^•^_2_^−^.

## Materials and methods

### Plant material Lilium longiflorum

Thunb anthers were collected from flowers and desiccated at 30 °C for 4 d; dry pollen was stored at −20 °C. Defrosted pollen samples were rehydrated in a humid chamber at 25 °C for 1 h. For EPR experiments tobacco pollen was used as well. Plants of *Nicotiana tabacum* L. var. Petit Havana SR1 were grown in a climatic chamber under controlled conditions (25°C, 16h light). Defrosted pollen was rehydrated in a humid chamber at 25 °C for 2 h.

### Pollen germination

For microscopy experiments lily pollen was mounted in a thin layer of 1,5% low-melting agarose (Sigma) on a cover glass and incubated in a small Petri dish in pollen germination medium (GM) at 29 °C for 60-120 min to achieve pollen tube initials, short or plng pollen tubes.

For EPR experiments lily or tobacco pollen was incubated in liquid GM at pollen concentration 2 mg/ml at 29 °C for 60 min. GM contained: 0.3 M sucrose, 1.6 mM H_3_BO_3_, 3 mM Ca(NO_3_)_2_, 0.8 mM MgSO_4_, 1 mM KNO_3_ in 50 mM MES-Tris buffer, pH 5.5 (lily) or 5.8 (tobacco).

### Exogenous ROS production

As a source of O_2_^•–^ we used riboflavin irradiated with UV (Beauchamp and Fridovich 1971; Smirnova et al. 2009). The freshly prepared solution of riboflavin (8 μM) in 50 mM K_2_HPO_4_ – KH_2_PO_4_ buffer (pH 7.4), containing 1mM EDTA, was irradiated with UV (365±12 nm) for 1 min. Controls for the effects of riboflavin on pollen tube growth have been performed and showed no effect.

Hydroxyl radical was generated in a Fenton reaction by adding 100 (membrane potential) or 500 (elsewhere) µM H_2_O_2_ and 100 µM FeSO_4_ (Kukavica et al. 2009) to pollen suspension.

### Intracellular ROS detection

Mitochondrial superoxide radical (O^•^_2_^−^) was detected with MitoSOX red (Thermo Fisher Scientific) (5 µM). Hydrogen peroxide was detected with pentafluorobenzenesulfonyl fluorescein (Santa Cruz Biotechnology) (10 µM). Staining was performed at 25°C for 10 min.

### Cytological analysis

Transport vesicles were identified by staining pollen tubes with (N-(3-triethylammoniumpropyl)-4-(6-(4-(diethylamino)phenyl)hexatrienyl) pyridinium dibromide; Molecular Probes) (8 µM). In the experiments testing the tube capacity for endocytosis, FM4-64 was present in incubation medium in the last 20 min of culture.

Mitochondria were identified by staining the pollen tubes with NAO (10-N nonyl-acridine orange; Molecular Probes) (5 µM).

### Membrane potential assessment

Mapping of MP in pollen tubes was performed as described earlier (Maksimov et al. 2018) with Di-4-ANEPPS (Sigma) which allows to detect accurately spatiotemporal variations of MP. Pollen tubes were stained with 2 μM Di-4-ANEPPS and immediately used for microscopy. Red fluorescence (>590 nm) was excited in two channels: blue (450–490 nm, F_b_) and green (540–552 nm, F_g_), F_b_/ F_g_ ratio is inversely proportional to MP value (higher ratio = depolarization).

Absolute MP values were determined with DiBAC_4_(3) (Bis-(1,3-dibutylbarbituric acid) trimethine oxonol, Molecular probes) which belongs to the group of slow dyes; charged molecules are distributed between the cell cytoplasm and surrounding medium in correspondence with the Nernst’s equation (Plášek and Sigler 1996). By measuring the fluorescence intensity of living (I) and fixed (completely depolarized) cells (I_0_) stained simultaneously with the same dye solution, membrane potential values (mV) are calculated according to the equation (Emri et al. 1998; Breygina et al. 2010):

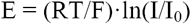

Cells were stained with the dye solution (5 µM) for 10 min. Measurements were performed in the apical domain.

### Assay of cytoplasmic pH

For pH mapping tubes were stained with 10 μM BCECF-AM (Invitrogen) at 25°C for 10 min. For pH measurement green fluorescence (515-565 nm) was excited in two channels: I_1_=475-495 nm; I_2_=426-446 nm. I_1_/I_2_ is proportional to pH value. Calibration both with nigericin-treated tubes and pseudocytosol was performed according to (Feijó et al. 1999).

### Fluorescent microscopy and computer image analysis

Localization of ROS and cytological analysis was performed with a confocal Nikon A1 microscope at the White Sea Biological Station of Moscow State University (Karelia, Russia) with 488 and 514 nm lasers.

Quantitative fluorescent microscopy was performed in a widefield fluorescence microscope Axioplan 2 imaging MOT (Zeiss) equipped with AxioCam HRc digital camera and a mercury lamp. For di-4-ANEPPS two filter sets were used: 450-490/>515 (blue excitation, Fb) and 540-552/>590 (green excitation, Fg). Filter change, shutter and camera are operated with AxioVision 4.7.10 (Zeiss) and obtained images were processed with the same software to assess Fb/Fg ratio in each point and perform the mapping; automatic program constructed for each type of acquisition experiment within AxioVision included filter sequence and standard exposure for each channel.

### EPR spectroscopy

The ability of pollen to remove OH• (generated in a Fenton reaction) was analyzed in experiments with spin trap DMPO (5,5-dimethyl-1-pylloline-N-oxide) (Sigma). First H_2_O_2_ and spin trap were added to a suspension of pollen, after 30 s the reaction was started by the addition of FeSO_4_. The spectra were recorded after 3.5 min of reaction at room temperature with RE-1307 spectrometer (Russia) operating at microwave power 20 mW, frequency 9.5 GHz, 1 min of sweep time. A manganese signal was used for standardizing the external signal.

### Statistical analysis

Average curves and meanings are presented in the figures; number of tubes used for each average is mentioned in the legends. Standard errors are shown for critical points of gradients. Significant difference was evaluated according to Student’s t test (*P < 0.05, **P < 0.01) and is shown on diagrams or mentioned in the text.

## Results

### ROS are unevenly distributed in lily pollen tube

We studied subcellular localization of ROS with confocal microscopy using pentafluorobenzenesulfonyl fluorescein (PFBSF) to visualize H_2_O_2_ (Maeda et al. 2004), and MitoSOX for mitochondria-derived superoxide-radical (Robinson et al. 2008). H_2_O_2_ is mostly seen in the cytosol and large spherical organelles (presumably, plastids) (Fig. 1a,d,g). The level of cytosolic H_2_O_2_ and the number of plastids can differ in individual tubes, but the pattern is stable: in most of the tubes H_2_O_2_ is concentrated in the apical area (Fig. 1a,d), but we have seen a number of tubes with H_2_O_2_ level uniformly high along the tube up to 50 µm from the tip (Fig. 1g). Mitochondria producing O^•^2^−^ were localized along the tube except the apical dome (Fig. 1b,e,h). We found that in pollen tubes H_2_O_2_ is poorly colocalized with mitochondrial O^•^_2_^−^ thus revealing mitochondria as a minor source of cytosolic H O (Fig. 1c,f,i).

**Fig. 1.**
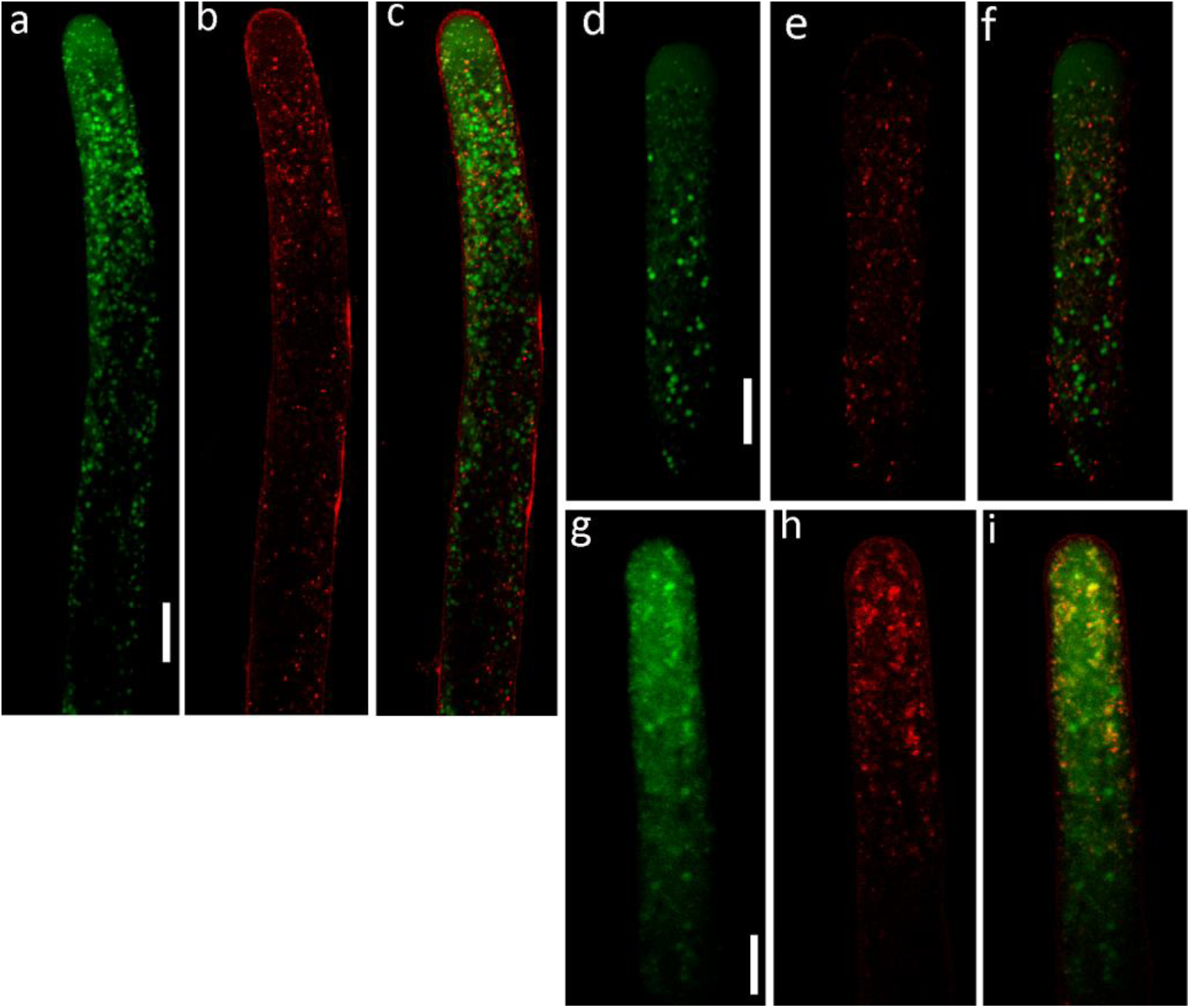
Localization of H_2_O_2_ and mitochondrial O^•^_2_^−^ in lily pollen tubes. Three randomly picked pollen tubes after 2 hours of incubation stained with pentafluorobenzenesulfonyl fluorescein (PFBSF) (a, d, g) and MitoSOX (b, e, h). c, f, i show colocalization of the two ROS. The tube in (c) has a higher level of ROS than the one in (f) (individual differences between cells), but the localization is the same: mitochondria producing O^•^_2_^−^ are accumulated in subapical part, H_2_O_2_ is poorly colocalized with mitochondria, but significant levels are as well detected in the apical cytoplasm and in plastids. The tube in (i) has the highest ROS level, and apical accumulation of H_2_O_2_ is not expressed. Bar – 20 µm

We used PFBSF to find out whether the apical accumulation of H_2_O_2_ is established early during the onset of polar growth. In pollen tube initials as well as in very short tubes an apical part of the tube is clearly stained marking H_2_O_2_-rich zone (Fig. 2a,d,f). In short tubes mitochondria-derived superoxide-radical can be stained as well, showing poor colocalization of H_2_O_2_ with mitochondria (Fig. 2b,c). Thus, in lily pollen the two endogenous ROS are unevenly and differentially distributed, probably, due to the separate production.

**Fig. 2.**
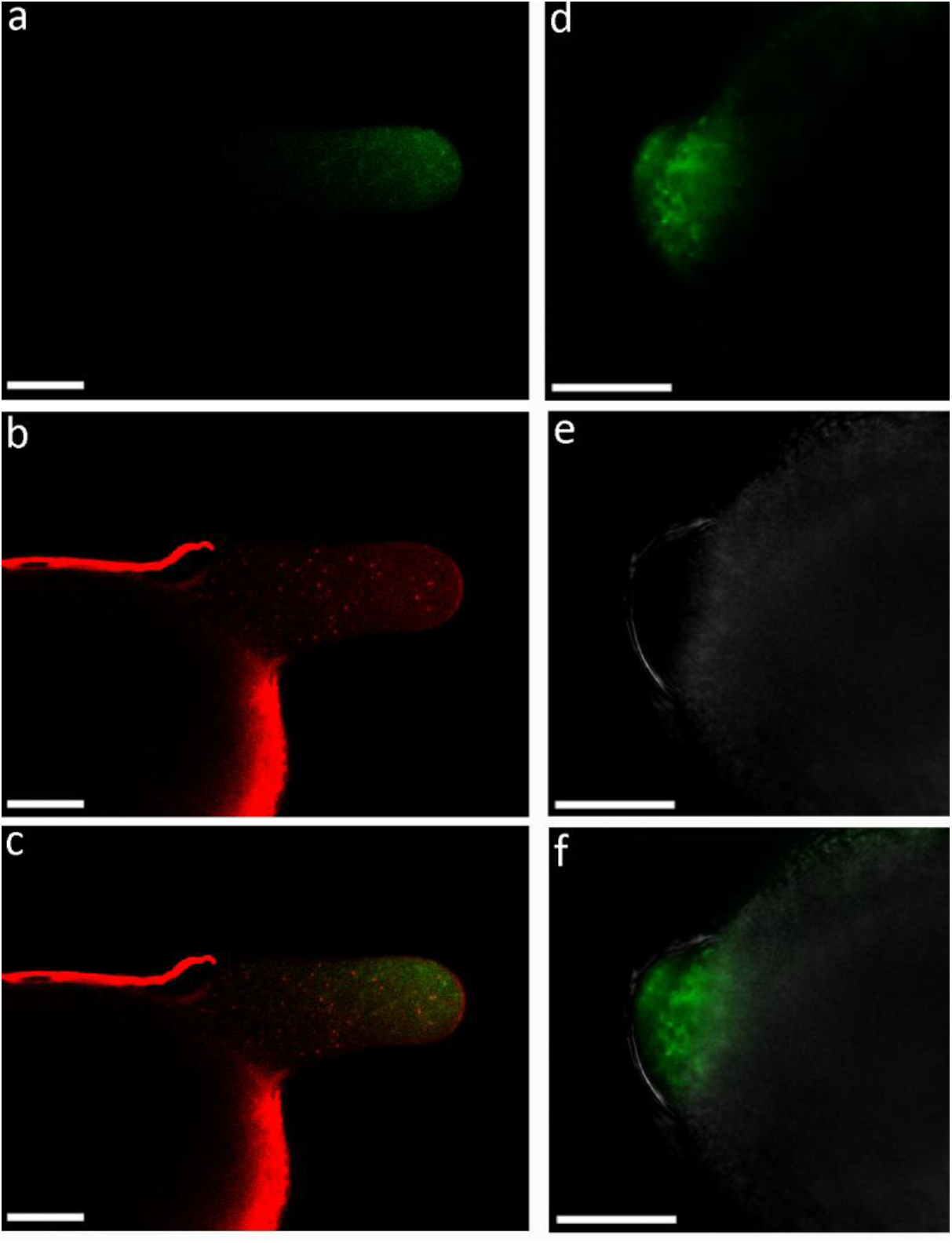
Localization of ROS in lily pollen tube initials and short pollen tubes. Typical short pollen tube (a-c) and pollen tube initial (d-f) stained with PFBSF (a, d, f) and MitoSOX (b). PFBSF demonstrates apical H_2_O_2_ accumulation in the cytoplasm of both structures; (c) shows poor colocalization of H_2_O_2_ with mitochondrial O^•^_2_^−^ in a short pollen tube. Pollen tube initials do not perform MitoSOX staining due to the small number of mitochondria; e shows corresponding DIC image. Bar – 20 µm

### ROS stimulate or inhibit pollen tube growth

To examine which ROS are the most important for pollen tube growth, including endogenous and those produced in the environment, we used different compounds affecting ROS formation/elimination balance: Mn-TMPP, which eliminates endogenous O^•^_2_^−^ and H_2_O_2_, exogenous H_2_O_2_, O^•^_2_^−^ (riboflavin/UV light), OH• (H_2_O_2_/FeSO_4_). All substances were present in the medium for the last 15 min of incubation period. Different effects of these reagents were found (Fig. 3a). OH• and MnTMPP decreased the speed of pollen tube growth, while O^•^_2_^−^ enhanced it. H_2_O_2_ (0.1 mM) had no effect on this value. Tubes under the influence of O^•^_2_^−^ were not only longer, but also thinner than in control (Fig. 3b). More than a half of pollen tubes growing in the presence of OH• and MnTMPP formed, on the contrary, expanded tips (tip 1.1 – 1.5 times wider than the shank) or ballooned tips (tip more than 1.5 times wider than the shank) (Fig. 3d,e): in both suspension only about 35% of tubes had normal shape after 15-min treatment compared to 92% in control (Fig. 3c). Thus, different ROS can be considered as accelerating or decelerating influences on pollen tube growth that possibly act *in vivo* as positive or negative signals controlling pollen physiology.

**Fig. 3.**
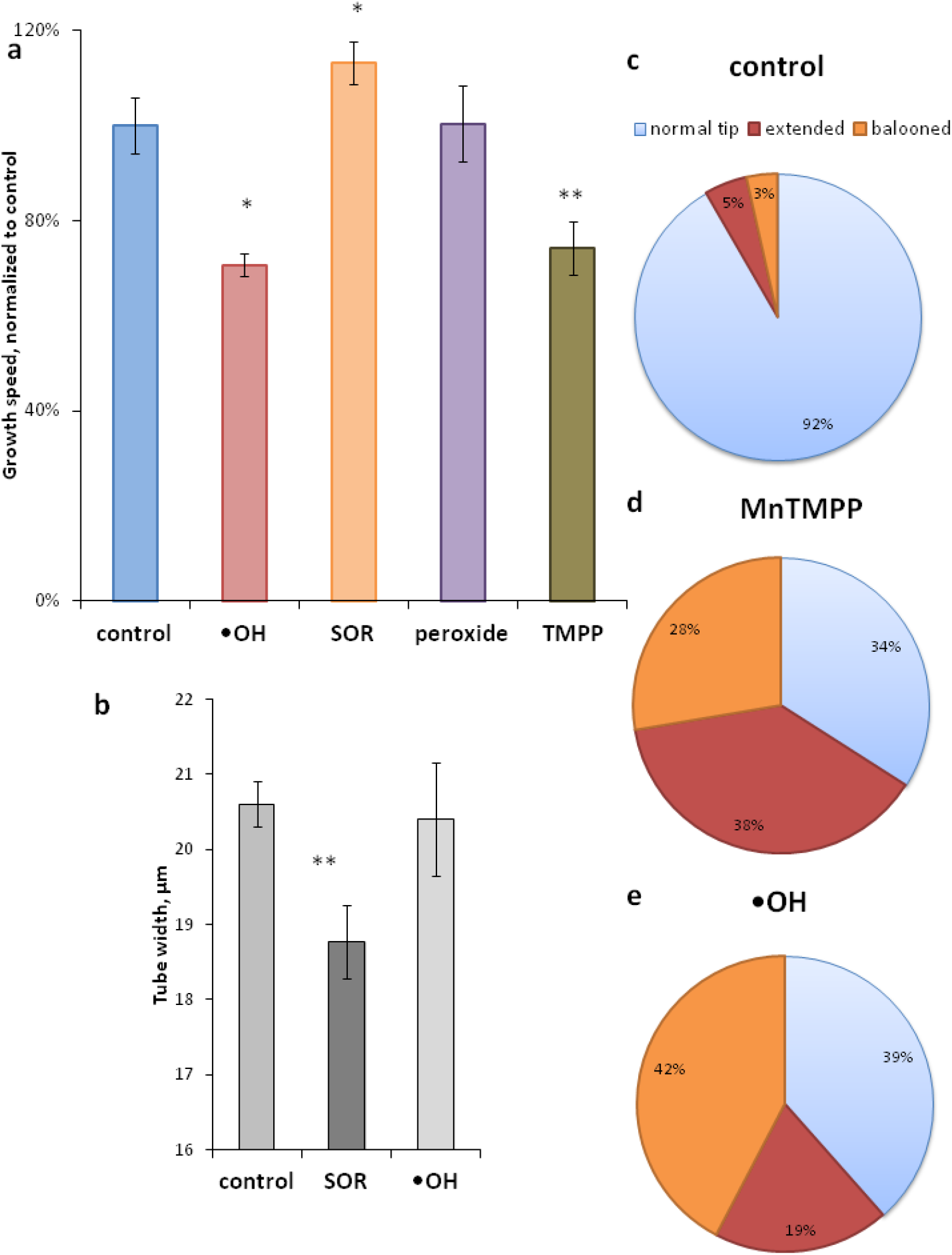
The effect of different ROS on pollen germination. a – pollen tube growth speed in control, in the presence of Mn-TMPP, which eliminates endogenous O^•^_2_^−^ and H_2_O_2_, exogenous H_2_O_2_, O^•^_2_^−^ (riboflavin/UV light) and OH• (H_2_O_2_/FeSO_4_) (absolute meanings normalized to control). H_2_O_2_ (100 µM) has no effect on growth speed, OH• and Mn-TMPP inhibit growth, O^•^_2_^−^ stimulates growth of pollen tubes. b – Mean pollen tube width. In control and O^•^_2_^−^-treated suspensions all tubes were measured, which revealed the “slimming” effect of SOR; in OH•-treated suspensions only non-extended parts of the tubes were measured; extended and ballooned tips will be discussed in (c). c-e – Mn-TMPP and OH• both affect the shape of the tube apex compared to control (c), but in TMPP-treated suspensions the most abundant pattern is “extended tip” (d), in OH•-treated – “ballooned tip” (e)

### ROS affect cytoplasm zoning

We found that seemingly opposite influences (addition of OH• and ROS quenching) have similar effects on pollen tube growth. But are these effects really the same? Cytological analysis provided an insight into these disturbances of polar growth. We used lipophylic dye FM4-64 to assess membrane transport (endo-/exocytosis) and NAO (binding to cardiolipin) to stain mitochondria and visualize their distribution in pollen tubes. Typical pattern of staining by both dyes is presented in Fig. 4a-c (control): FM4-64 staining plasmalemma along the shank and an apical cone of exocytic vesicles, mitochondria are located all along the tube length, except the apical dome, with less density in the apical part. In the presence of Mn-TMPP the apical zone of the tube is expanded and contains multiple mitochondria, thus less of them enter the “apical dome” (Fig. 4e). The apical accumulation of vesicles is even more pronounced than in control (Fig. 4d), demonstrating active membrane transport and exocytosis, which probably causes the tip extension. In total, organelle distribution is not impaired in these tubes (Fig. 4f). In OH•-treated tubes the tip is also expanded with mitochondria evenly distributed along the tube (Fig. 4i). FM staining reveales severe disturbances of membrane transport in all OH•-treated tubes: only plasma membrane is stained (Fig. 4h). No internalization and/or recycling of the dye occur. Cytoplasm zoning is completely absent (Fig. 4g).

**Fig. 4.**
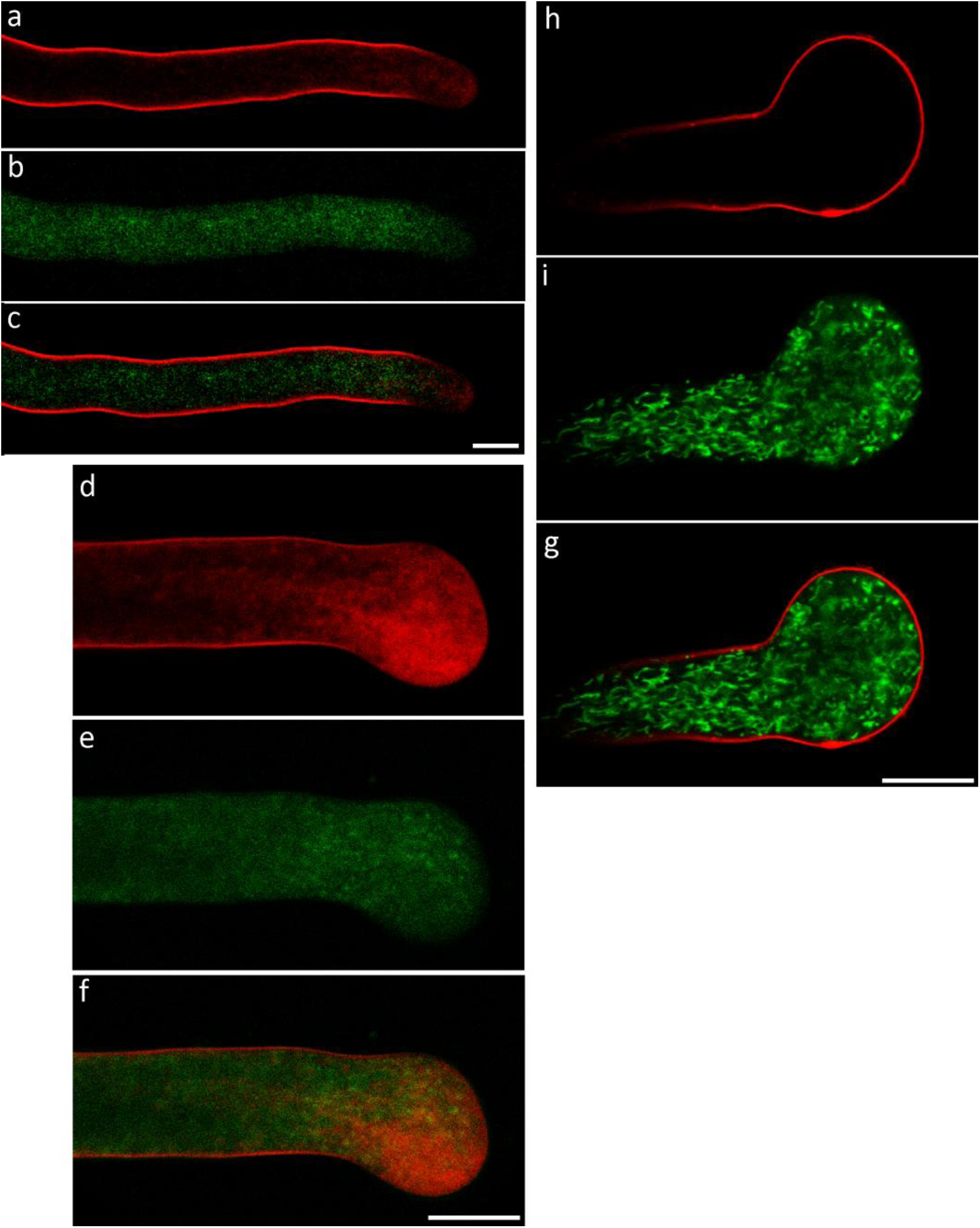
Organelle distribution in pollen tubes with impared redox-status. Pollen tubes after 2 hours of incubation stained with FM4-64 for membrane material (a, d, h) and NAO for mitochondria (b, e, i); c, f, g shows colocalization of organelles. Control tube (a-c) shows typical staining with vesicles in the apical area and mitochondria distributed along the tube. Mn-TMPP (15 min) affects shape of the apex, membrane traffic is active (d), mitochondria are less abundant in the apical area (e), thus total pattern of organelle distribution is unaffected (f). OH• completely blocks vesicle transport (h), mitochondria are evenly distributed throughout the tube including ballooned apex (i). Colocalization demonstrates the loss of polarity (g). Bar – 20 µm

### Oxygen radicals differentially shift membrane potential values

One of the most important values reflecting pollen tube zoning is membrane potential (MP), which forms longitudinal gradient that has been seen to date in all species studied including, recently, lily pollen tubes (Podolyan et al. 2019). We used two independent optical methods to assess MP level: slow cytoplasmic dye DiBAC_4_(3) to reveal shifts of absolute MP values in the apical area of the tube and fast membrane-located dye Di-4-ANEPPS – to study the effect on MP gradient along the pollen tube. DiBAC_4_(3) demonstrated opposite effects of radicals on MP: OH• caused depolarization while O^•^_2_^−^ caused hyperpolarization of pollen tube membrane (Fig. 5a). These effects correlated with growth speed in both cases. For OH• only non-balooned tubes were measured, as ballooned tips stain differently, which can cause artifacts. Still, the effect was significant.

**Fig. 5.**
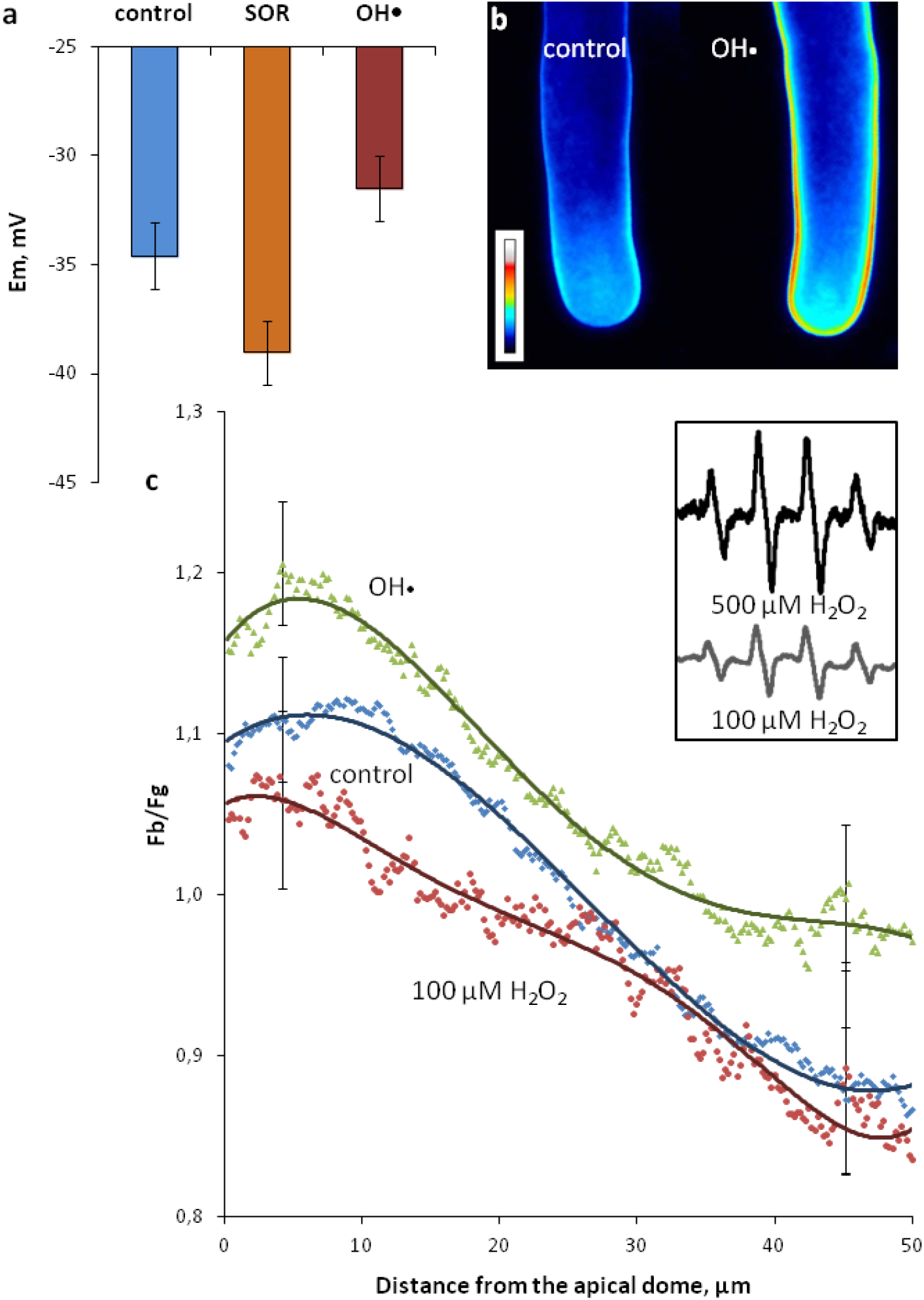
Membrane potential in lily pollen tubes assessed by ratiometric Di-4-ANEPPS staining and DiBAC_4_(3) cytoplasmic staining. a – O^•^_2_^−^ and OH• cause opposite effects on membrane potential in lily pollen tubes (apical part): significant hyperpolarization is found in SOR-treated tubes compared to control assessed by DiBAC_4_(3) cytoplasmic staining; OH• causes depolarization; b, c – in control tubes there is a lateral gradient: subapical zone is significantly depolarized compared to distal zone (Di-4-ANEPPS membrane staining). b – typical MP-sensitive staining of pollen tubes, LUTs were applied after image division (fluorescence induced by blue light to that induced by green light). c – Average curves from more than 18 pollen tubes. 100 µM H_2_O_2_ has no effect on the gradient, the same concentration of H_2_O_2_ after the addition of FeSO_4_ and subsequent OH• production causes significant depolarization in all regions of the tubes. Inset – the level of OH• production assessed by EPR spectroscopy with DMPO spin trap

Using Di-4-ANEPPS we also studied the effect OH• on MP gradient: depolarization occurred in proximal subapical and distal parts of the tubes as well as in the apical part (Fig. 5b,c). For O^•^_2_^−^ unfortunately it was impossible to assess MP gradient shifts, as riboflavin fluorescence interferes with the dye. Hyperpolarization found with DiBAC_4_(3) was checked not to be caused by riboflavin itself.

To produce OH• in this experiment we used the mixture of 100 µM H_2_O_2_ and FeSO_4_. H_2_O_2_ is well known to affect MP in pollen tubes (Maksimov et al. 2018; Podolyan et al. 2019), so, to check if the depolarization is caused by H_2_O_2_ itself, we added 100 µM H_2_O_2_ without FeSO_4_. No depolarization occurred, specifying that the effect is of OH•. We did not use higher radical concentration, so that the effects of H_2_O_2_ and OH• would not interfere, as 500 µM H_2_O_2_ causes membrane hyperpolarization (Podolyan et al. 2019). To check relative quantity of OH• produced from 100 and 500 µM H_2_O_2_ we used EPR-spectroscopy with DMPO spin trap (Fig. 5, inset).

### Superoxide radical affects cytoplasmic pH

Important value connected to cytoplasm zoning is intracellular pH. In control pollen tubes typical pH gradient can be seen with acidic tip, alkaline subapical zone (“band”) and nearly neutral distal zone (Fig. 6). Superoxide radical causes alkalinization of the tube; in the apical zone (1 µm from the tip) it is less pronounced while in subapical (10 µm) and distal (50 and 90 µm) there is a significant increase of pH values (P<0,95). The shape of the gradient remains similar.

**Fig. 6.**
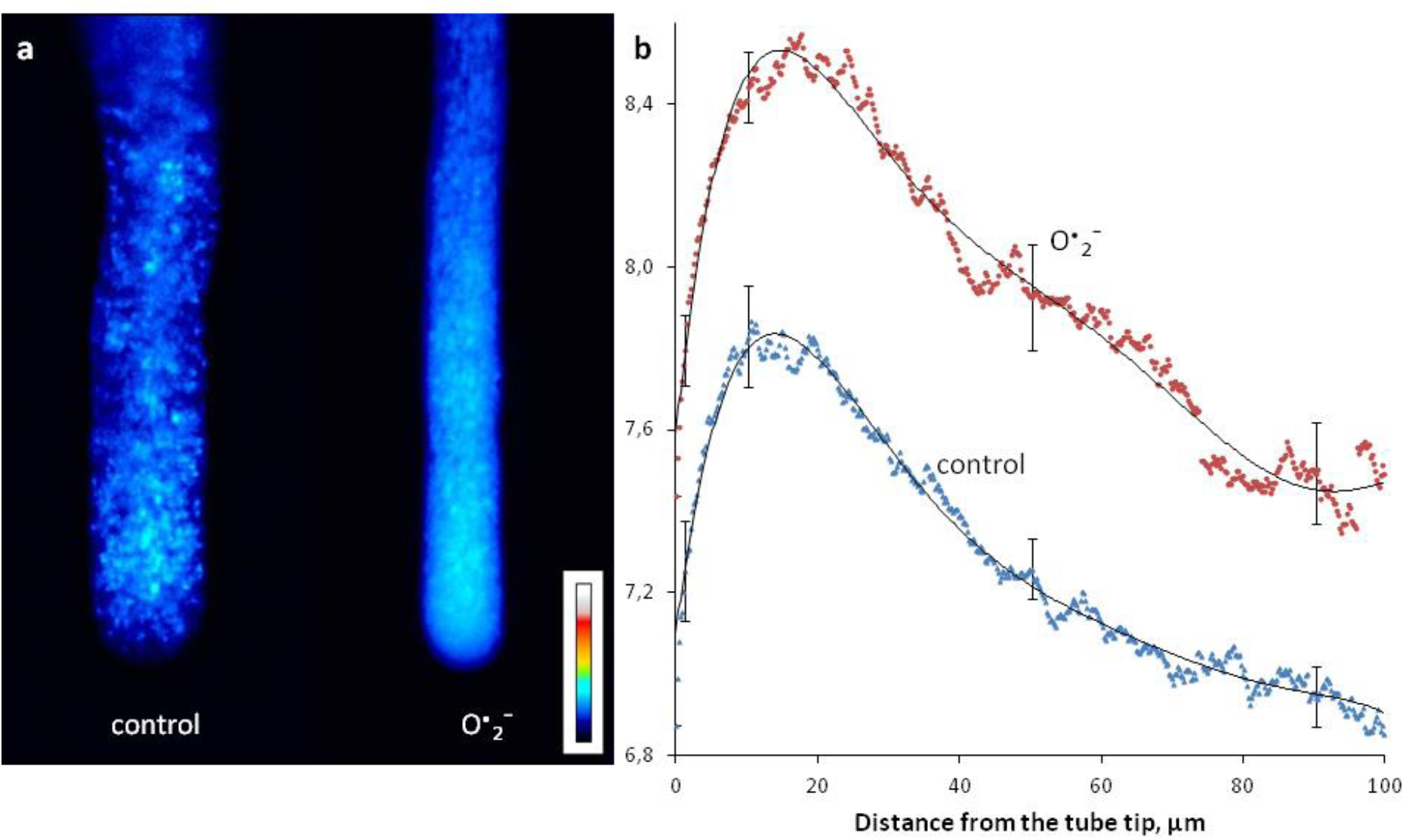
Cytoplasmic pH gradient affected by O^•^_2_^−^ assessed by ratiometric BCECF-AM staining. In control tubes a typical gradient can be seen: apical dome has pH below 7, “alkaline band” is localized in subapical part, and distal part of the tube is rather neutral. O^•^_2_^−^ (riboflavin/UV light) has no significant effect on the shape of the gradient, but causes significant alkalization with the most noticeable effect in subapical and distal regions. a – typical pH-sensitive staining of pollen tubes, LUTs were applied after image division (fluorescence in pH-sensitive channel to that in pH-insensitive). b – Average curves from more than 25 pollen tubes

### OH• is quenched by pollen

We used EPR spectroscopy to evaluate the radical content in pollen suspension with DMPO spin trap. We found that both lily and tobacco pollen significantly reduced DMPO signal in all experiments where Fenton reaction was performed (Fig. 7), thus demonstrating antioxidant activity of both pollen suspensions. Besides, we checked, whether OH• is produced from 1 mM H_2_O_2_ in the presence of lily pollen. We found that no DMPO signal can be detected without the addition of FeSO_4_. Thus, pollen can quench but not produce OH• in detectable amounts.

**Fig. 7.**
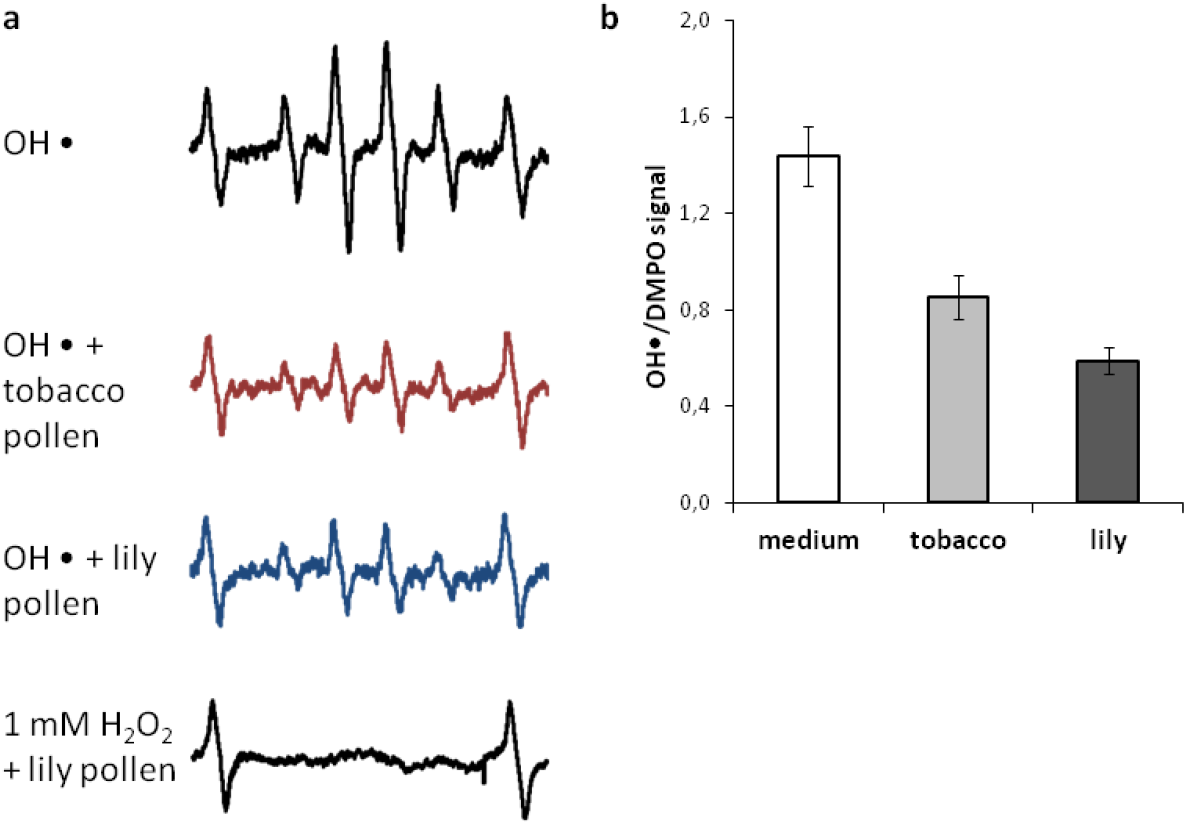
Level of OH• in germination medium and pollen suspension measured by EPR spectroscopy. a – typical EPR signals from OH• adducts obtained with DMPO spin trap and the outer two peaks correspond to the Mn^2+^ markers. Relatively high level of signal can be seen in pollen-free medium in the presence of 500 µM H_2_O_2_/FeSO_4_; both lily and tobacco pollen reduce the signal, demonstrating antioxidant properties. In the presence of pollen no measurable amounts of OH• are produced from H_2_O_2_ (without FeSO_4_). b – Average EPR signals from OH• adducts obtained with DMPO spin trap. All signals are normalized using the amplitudes of the Mn^2+^ signal.

## Discussion

To date convincing evidences for the involvement of H_2_O_2_ in controlling polar growth has been obtained and acknowledged (Maksimov et al. 2016, 2018), but the role for other ROS was unclear as they were usually considered all together as “total ROS” (Bell et al. 2009; Mangano et al. 2016; Feijó and Wudick 2018). We used fast growing lily pollen tubes as a model system to check the possible involvement of radical ROS in the regulation of polar growth. Produced from riboflavin activated with UV light in germination medium, after 10 min of incubation O^•^_2_^−^ caused growth stimulation, tube thinning, cytoplasm alkalinization and plasma membrane hyperpolarization. Theoretically, this effects could be attributed to H_2_O_2_ which can be produced from O^•^_2_^−^ by SOD or in a non-enzymatic reaction (Mangano et al. 2016). Earlier hyperpolarization had been found in tobacco pollen protoplasts (Maksimov et al. 2016) and growth stimulation was found in *Picea pungens* pollen tubes (Maksimov et al. 2018). Here we checked these effects and did not find any of them in the presence of 0.1 mM H_2_O_2_ which is the effective concentration modulating ion currents in lily pollen protoplasts (Breygina et al. 2016). The minimal concentration of H_2_O_2_ to cause hyperpolarization in lily pollen tubes was 0.5 mM, to provoke pH shift – 1 mM (Podolyan et al. 2019). Thus, both O^•^_2_^−^ and H_2_O_2_ can be considered as “stimulating” ROS, but the effects of O^•^_2_^−^ are more pronounced. O^•^_2_^−^ is the primary ROS to be produced by NADPH-oxidase in pollen (Potocký et al. 2012; Jimenez-Quesada et al. 2019) and, probably, in female tissues during pollination (McInnis et al. 2006; Zafra et al. 2010; Duan et al. 2014), but O^•^_2_^−^ cannot spread over noticeable distances, unlike H_2_O_2_, thus both ROS can activate polar growth and cause other physiological effects.

Unlike O^•^_2_^−^, OH• severely inhibits growth, causes membrane depolarization in different tube zones and affects cytoplasm organization. More than 60% of pollen tubes form expanded or ballooned tips. This effects agrees well with previously postulated hypothesis on the involvement of the radical in the regulation of the mechanical properties of the cell wall during early stages of pollen germination (Smirnova et al. 2013): in this work OH• was found to loosen pollen intine and reduce the resistance of pollen to hypotonic stress. Besides the effect on the cell shape, in OH•-treated pollen tubes we also found blocked membrane trafficking and membrane depolarization. Previously similar shifts of membrane potential gradient were found in tobacco (Breygina et al. 2009) and spruce (Maksimov et al. 2018) pollen tubes exposed to H^+^-ATPase inhibitor orthovanadate. Inhibition of this enzyme by OH• could lead to the observed effects. Previously effects of oxygen radicals on this enzyme have not been studied thoroughly, but there is data on the involvement of OH• in Fe^2+^-induced inhibition of H^+^-ATPase (Yang et al. 2003).

Here we used relatively low concentration of Fenton reagents compared to those routinely applied to plant cells (Demidchik et al. 2010; Smirnova et al. 2013); besides, pollen tubes significantly reduced OH• level in the incubation medium, thus demonstrating antioxidant properties. Antioxidant properties had been previously reported for tobacco: pollen diffusates (Žárský et al. 1987) interfered with the standard guaiacol peroxidase assay while pollen exine (Smirnova et al. 2012) reduced the level of stable nitroxyl radical TEMPO and DMPO/OH• adduct obtained in a Fenton reaction. Negative effects of hydroxyl radical found here despite low level of OH• in germination medium suggest that this radical can inhibit and disrupt pollen tube growth in the pistil not due to total oxidative stress, but via specific targets. To date selective sensitivity to OH• has been found for cation channels in root cells (Demidchik 2003; Demidchik et al. 2010).

The importance of endogenous ROS produced by NOX is demonstrated by the inhibition of growth by superoxide dismutase /catalase mimic MnTMPP which was earlier found to inhibit pollen germination in kiwifruit (Speranza et al. 2012), tobacco (Smirnova et al. 2013) and spruce (Maksimov et al. 2018). Effect of MnTMPP, which quenches both H_2_O_2_ and O^•^_2_^−^, was close in these cases to effects of DPI as both reagents block the production of endogenous ROS. Moreover, here we demonstrate that MnTMPP causes cytoplasm reorganization and subsequent alteration of cell shape. This can be caused by the violation of calcium gradient or pH gradient in the cytoplasm which have been found in these tubes (Podolyan et al. 2019) and subsequent alterations of actin structure. However, vesicle traffic in ballooned pollen tubes in the presence of MnTMPP is very active, as demonstrated by FM staining.

The spatial distribution of endogenous ROS in pollen tubes revealed both apical accumulation of H_2_O_2_ probably formed from NOX-derived O^•^_2_^−^ and produced by polyamine oxidase (Wu et al. 2010), and significant signal in plastids along the shank. This pattern is quite similar to the one found in *Picea* pollen tubes (Maksimov et al. 2018), but, as expected, apical accumulation of H_2_O_2_ is larger in *Lilum*, with higher growth speed. Apical H_2_O_2_ accumulation was as well found very short pollen tubes which agrees well with the pattern reported recently for *Arabidopsis* (Do et al. 2019). Localization of H_2_O_2_ in plastids is similar in lily and spruce, though they are much larger in *Picea* and contain a lot of starch (Breygina et al. 2019). Nonetheless, in angiosperms plastids are as well the largest organelles in pollen tubes (Southvvorth and Dickinson 1981; Staff et al. 1989; Fujiwara et al. 2012). No significant colocalization of H_2_O_2_ with mitochondria-derived O^•^_2_^−^ was found, which contradicts the previously made assumption that mitochondria which are the main source of ROS (Cárdenas et al. 2006), at least for certain species.

## Declarations

### Ethics approval

Not applicable

### Consent to participate (include appropriate statements)

Not applicable

### Consent for publication (include appropriate statements)

Not applicable

### Availability of data and material (data transparency)

Raw data can be provided upon request

### Code availability

Not applicable

### Authors’ contributions

MB conceived and designed research, AP and OL conducted experiments. AP and MB analyzed data. MB wrote the manuscript. All authors read and approved the manuscript.

